# Pathogenicity Prediction of GABA_A_ Receptor Missense Variants

**DOI:** 10.1101/2023.11.14.567135

**Authors:** Ya-Juan Wang, Giang H. Vu, Ting-Wei Mu

## Abstract

Variants in the genes encoding the subunits of gamma-aminobutyric acid type A (GABA_A_) receptors are associated with epilepsy. To date, over 1000 clinical variants have been identified in these genes. However, the majority of these variants lack functional studies and their clinical significance is uncertain although accumulating evidence indicates that proteostasis deficiency is the major disease-causing mechanism for GABA_A_ receptor variants. Here, we apply two state-of-the-art modeling tools, namely AlphaMissense, which uses an artificial intelligence-based approach based on AlphaFold structures, and Rhapsody, which integrates sequence evolution and known structure-based data, to predict the pathogenicity of saturating missense variants in genes that encode the major subunits of GABA_A_ receptors in the central nervous system, including *GABRA1*, *GABRB2*, *GABRB3*, and *GABRG2*. Our results demonstrate that the predicted pathogenicity correlates well between AlphaMissense and Rhapsody although AlphaMissense tends to generate higher pathogenic probability. Furthermore, almost all annotated pathogenic variants in the ClinVar clinical database are successfully identified from the prediction, whereas uncertain variants from ClinVar partially due to the lack of experimental data are differentiated into different pathogenicity groups. The pathogenicity prediction of GABA_A_ receptor missense variants provides a resource to the community as well as guidance for future experimental and clinical investigations.

## INTRODUCTION

Epilepsy is one of the most common neurological diseases in the world with a broad phenotypic spectrum ^[1]^. Recent advances in genome sequencing identified an increasing number of genes that are associated with epilepsy ^[2]^. According to protein functions, epilepsy-associated genes can be grouped to ion channels, enzymes and enzyme modulators, transports and receptors, and others ^[3]^. Genetic epilepsy is often linked to developmental delay, movement disorder, and other comorbidities ^[4]^. Due to the important role of neurotransmitter-gated ion channels in controlling the excitation-inhibition balance in the central nervous system, genes encoding these ion channels, including excitatory N-methyl-D-aspartate (NMDA) receptors and inhibitory γ-aminobutyric acid type A (GABA_A_) receptors, are recognized as prominent epilepsy-causing genes ^[5]^. Here, we focus on GABA_A_ receptors, the primary inhibitory neurotransmitter-gated ion channels in the human brain ^[6]^. They mediate the fast inhibitory GABA-induced chloride currents and hyperpolarize the postsynaptic members to reduce neuronal firing.

Proteostasis maintenance of GABA_A_ receptors is essential for their function in the central nervous system ^[7]^. GABA_A_ receptors are assembled as pentamers from a specific combination of 19 subunits, including α1-α6 (GABRA1-A6), β1-β3 (GABRB1-B3), γ1-γ3 (GABRG1-G3), δ (GABRD), ε (GABRE), θ (GABRQ), π (GABRP), and ρ1-ρ3 (GABRR1-R3). The distribution of GABA_A_ receptors is throughout the brain regions, and the most abundant subtype is composed of two α1 subunits, two β2 subunits, and one γ2 subunit ^[8]^. To function, GABA_A_ receptor subunits need to fold in the endoplasmic reticulum (ER) with the assistance of molecular chaperones and subsequently assemble with other subunits to form heteropentamers. The properly assembled receptors exit the ER and traffic to the plasma membrane to act as chloride channels. Unassembled and misfolded subunits are retained in the ER, which could be routed to the degradation pathway by the ER-associated degradation ^[9]^. Recent quantitative proteomics analysis identified the proteostasis network that regulates the folding, assembly, trafficking, and degradation of GABA_A_ receptors ^[10]^.

Recent cryo□electron microscopy (cryo-EM) studies solved the high-resolution structures of pentameric GABA_A_ receptors, including α1β2γ2 receptors ^[11]^ and α1β3γ2 receptors ^[12]^. The pentameric receptors are arranged as β-α1-β-α1-γ2 counterclockwise when viewed from the synaptic cleft (**Figure 1A**). Each pentamer has two binding sites for the neurotransmitter, GABA, at the interfaces between β subunits and α1 subunits. Residues from β subunits constitute the principal binding site, denoted as “positive” (+) side, whereas residues from α1 subunits constitute the complementary binding site, denoted as “negative” (−) side. Each subunit shares a common structural scaffold, including a large extracellular N-terminal domain (NTD), four transmembrane helices (TM1-TM4), and loops connecting transmembrane helices (a short intracellular TM1-2 loop, a short extracellular TM2-3 loop, and a long intracellular TM3-4 loop), and a short extracellular C-terminus (**Figure 1B**, **1C**). The secondary structures of the NTD contain two α-helices, ten β-sheets (β1-β10), and connecting loops (**Figure 1C**, **1D**). GABA_A_ receptors belong to the Cys-loop receptor superfamily ^[7]^. The signature Cys-loop in GABA_A_ receptor subunits is designated as loop 7. Biochemical studies revealed that several segments in GABA_A_ receptor subunits play an important role in binding the ligand: the binding loops in the principal side are called loop A-C, whereas the binding loops in the complementary side are called loop D-F (**Figure 1C**, **1D**).

**Figure 1.**
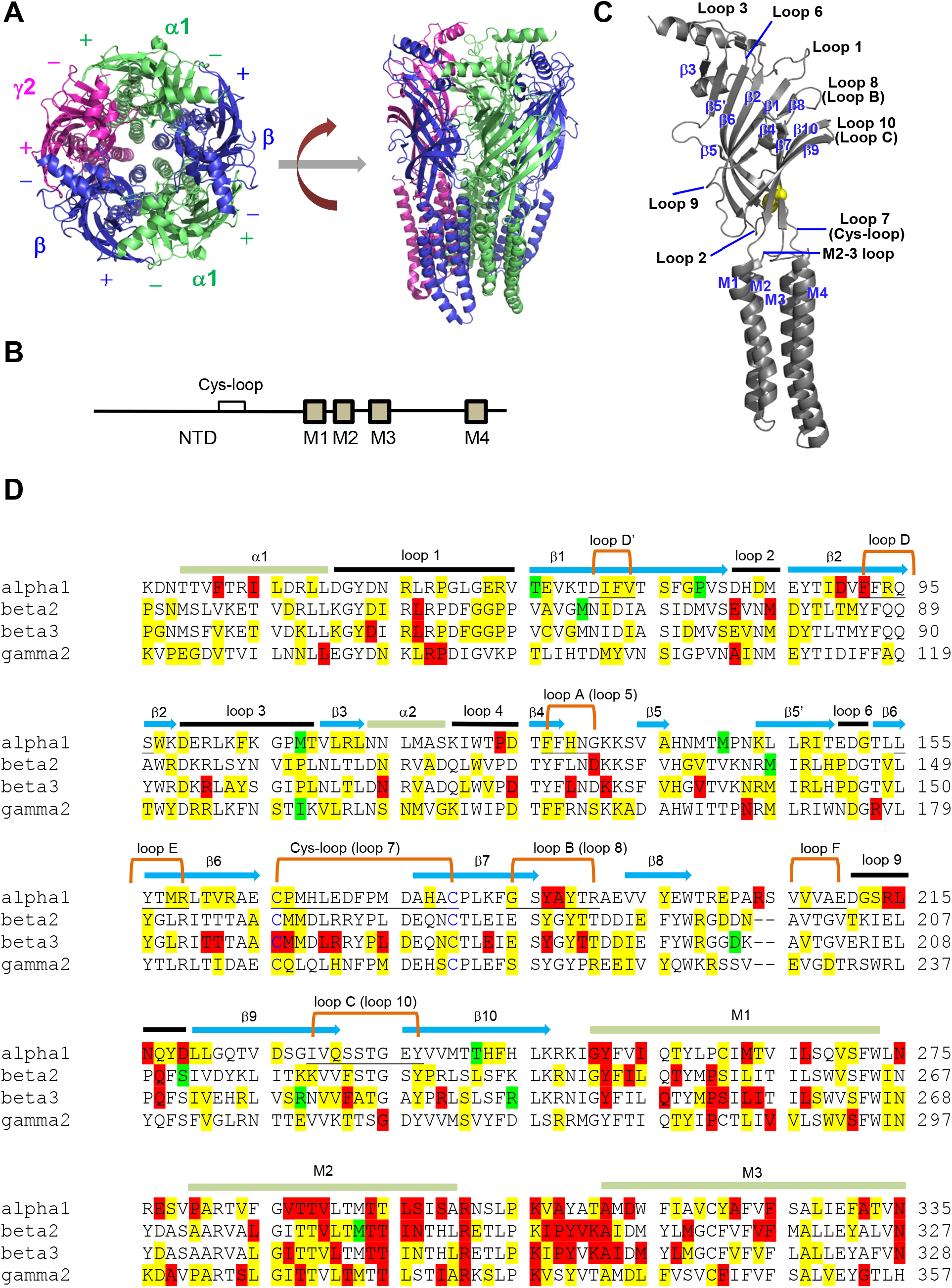
Structures and sequence alignment of GABA_A_ receptors. (**A**) Cartoon representation of pentameric α1βγ2 receptors, built from 6X3S.pdb. The principal side of one subunit is denoted as “+”, whereas the complementary side of one subunit is denoted as “−”. (**B**) The schematic of the primary protein sequence of a GABA_A_ receptor subunit. NTD, N-terminal domain; M1-M4, transmembrane helices 1 to 4. (**C**) The secondary structures of a GABA_A_ receptor subunit. The two cysteines in the signature Cys-loop are colored in yellow. (**D**) The sequence alignment of major human GABA_A_ receptor subunits, including α1, β2, β3, and γ2. The residue positions that harbor clinical missense variants are highlighted. According to ClinVar annotation, pathogenic variants are colored in red, uncertain in yellow, and benign in green.

To date, over 1000 clinical variants in genes encoding GABA_A_ receptor subunits have been recorded in ClinVar (www.clinvar.com), including missense, nonsense, and frameshift variants. However, the clinical significance of these variants is not adequately addressed since most of them lack functional characterization and many of them are classified as uncertain or conflicting interpretations. For the limited number of characterized GABA_A_ receptor variants, accumulating evidence indicated that proteostasis deficiency that resulted from misfolding and excessive degradation of the variants is a major disease-causing mechanism ^[13]^. Adapting the ER proteostasis network pharmacologically corrected the misfolding and restored the surface trafficking and thus ion channel function for a variety of pathogenic GABA_A_ receptor variants ^[14]^. Another important disease-causing mechanism is that the missense variants result in the channel gating defects and altered electrophysiological properties, such as current kinetics and current amplitude.

Here, we applied two state-of-the-art modeling tools, namely AlphaMissense ^[15]^ and Rhapsody ^[16]^, to comprehensively predict the pathogenicity of saturating missense variants in the major subunits of GABA_A_ receptors (α1, β2, β3, and γ2). Furthermore, we compared the prediction with the current ClinVar clinical data, aiming to provide guidance for future experimental investigation.

## RESULTS and DISCUSSION

### Distribution of clinical missense variants in ClinVar (CV) in the protein sequences of GABA_A_ receptor subunits

We first utilized the knowledge about the clinical significance of GABA_A_ variants according to ClinVar (CV) ^[17]^, which uses standard terms for Mendelian diseases recommended by the American College of Medical Genetics and Genomics (ACMG) and the Association for Molecular Pathology (AMP), including “pathogenic,” “likely pathogenic,” “uncertain significance,” “likely benign,” and “benign” ^[18]^. We simplified the interpretation of clinical variants to three classes: Class I, the pathogenic class, includes both “pathogenic” and “likely pathogenic”; Class II, the uncertain class, includes “uncertain significance” as well as “conflicting interpretations”; Class III, the benign class, includes both “likely benign,” and “benign”. The positions of all known clinical missense variants in the major subunits of GABA_A_ receptors, including GABRA1 (α1), GABRB2 (β2), GABRB3 (β3), and GABRG2 (γ2), were highlighted in their protein sequence alignment (**Figure 1D** and **Supplemental Figure S1**). Clearly, for a total of 889 missense variants, including 232 in GABRA1, 214 in GABRB2, 232 in GABRB3, and 211 in GABRG2, they are distributed throughout the primary protein sequence, including the signal peptide, NTD, TM1-4, and loops connecting transmembrane helices.

All clinical variants in TM1, TM2, and TM3 except one are interpreted as pathogenic or uncertain (**Figure 1D** and **Supplemental Tables S1, S2, S3, S4**), indicating that the first three transmembrane helices are prone to disease-causing variations. Variants in TM1 and TM3 were reported to lead to proteostasis defects due to protein misfolding and excessive degradation of the subunits carrying the variations, resulting in loss of function of GABA_A_ receptors ^[13b]^. TM2 lies in the innermost part of the central pore and plays a critical role in channel gating; also the amino acid sequences in TM2 are well conserved, (for example, the TTVLTMTT motif is conserved in all the four subunits); therefore, many variants in TM2 are classified as pathogenic. Indeed, electrophysiology experiments demonstrated that T287P and I288S in β2 and T287P and T288N in β3 reduced the peak current amplitudes ^[19]^. In sharp contrast, all the variants in TM4 are interpreted as benign or uncertain (**Supplemental Figure S1**), suggesting that being synthesized and folded at the late stage during the biogenesis step, variations in TM4 could be better tolerated to conserve the structure and function of GABA_A_ receptors compared to TM1- TM3. Also TM4 occupies the outermost side of the transmembrane domain.

Many variants in the ligand-binding pockets, especially in the principal binding loops (Loop A, Loop B (aka loop 8), and Loop C), are categorized as pathogenic (**Figure 1D**), consistent with the critical role of these loops in binding the ligands to induce the conformational change of the channels to open the ion pore. In addition, several connecting loops in the NTD (**Figure 1C**), such as loop 2 connecting β2 sheet and β1 sheet, loop 7 (aka Cys-loop) connecting β7 sheet and β6 sheet, and loop 9 connecting β9 sheet and β8 sheet, harbor many pathogenic variants (**Figure 1D**), suggesting that maintaining the structural integrity of these loops are important for the function of GABA_A_ receptors. It is also worth noting that the TM2-3 loop is a hot spot for pathogenic variants (**Figure 1D**), possibly because this loop couples the ligand-binding activity to the channel gating process and thus variants in this region would reduce the channel function substantially. In sharp contrast, almost all variants in the large cytosolic TM3-4 loop are interpreted as uncertain or benign (**Supplemental Figure S1**), suggesting that this region, being unstructured, is well tolerated for variations. Concomitantly, while determining the structures for pentameric GABA_A_ receptors, usually researchers replaced the large TM3-4 loop with a seven-amino acid linker ^[11b, 12, 20]^.

### AlphaMissense (AM) predictions of GABA_A_ receptor variant pathogenicity

The interpretation of clinical significance for GABA_A_ variants in ClinVar is inadequate because 644 among the known 889 missense clinical variants in the major subunits (168 among 232 variants in GABRA1, 141 among 214 variants in GABRB2, 162 among 232 variants in GABRB3, and 173 among 211 variants in GABRG2) are classified as “uncertain” or “conflicting interpretations”, indicating that 72% of these missense clinical variants need further investigation for their clinical significance. In addition, the clinical classification in ClinVar according to ACMG/AMP is subjected to human bias.

To fill the gap for the interpretation of missense variants, we used two state-of-the-art tools, namely AlphaMissense (AM) ^[15]^ and Rhapsody (RS) ^[16]^, to predict the pathogenicity of saturating amino acid substations in the major subunits of GABA_A_ receptors. AM makes pathogenicity predictions by combining structural context, which is achieved by AlphaFold ^[21]^, and evolutionary conservation ^[15]^. Pathogenicity classification cut-off values are calculated by AM. Although the cut-off values may be different for individual proteins, in the cases of GABRA1, GABRB2, GABRB3, and GABRG2, these GABA_A_ receptor subunits all share the same cut-offs as the following: scores < 0.340 as benign, scores from 0.340 to 0.564 as ambiguous, and scores > 0.564 as pathogenic.

We first evaluated the AM performance on clinical missense variants with the ClinVar annotations. The spatial distributions of the clinical variants were displayed in the three-dimensional cryo-EM structures of GABA_A_ receptor subunits: **Figure 2A**, **3A**, and **5A** showed the positions of α1, β2, and γ2 variants using 6X3S.pdb ^[11a]^, whereas **Figure 4A** showed the positions of β3 variants using 6HUK.pdb ^[22]^. The AM prediction correlated well with CV annotations for variants in Class I, the pathogenic class, and Class III, the benign class. In Class I, the pathogenic class, for GABRA1, CV had a total of 48 variants, among which AM predicts 42 (**Table 1**); for GABRB2, CV had a total of 39 variants, among which AM predicts 39 (**Table 2**); for GABRB3, CV had a total of 52 variants, among which AM predicts 51 (**Table 3**); for GABRG2, CV had a total of 26 variants, among which AM predicts 26 (**Table 4**). In Class III, the benign class, for GABRA1, CV had a total of 16 variants, among which AM predicts 9 (**Table 1**); for GABRB2, CV had a total of 34 variants, among which AM predicts 24 (**Table 2**); for GABRB3, CV had a total of 18 variants, among which AM predicts 18 (**Table 3**); for GABRG2, CV had a total of 12 variants, among which AM predicts 12 (**Table 4**). The major difference between AM prediction and CV annotation was within Class II, the uncertain class. AM prediction can differentiate the large amount of CV uncertain variants to pathogenic, uncertain, and benign (**Supplemental Tables S1-S4**), providing valuable guidance for future investigation (also see below).

**Figure 2.**
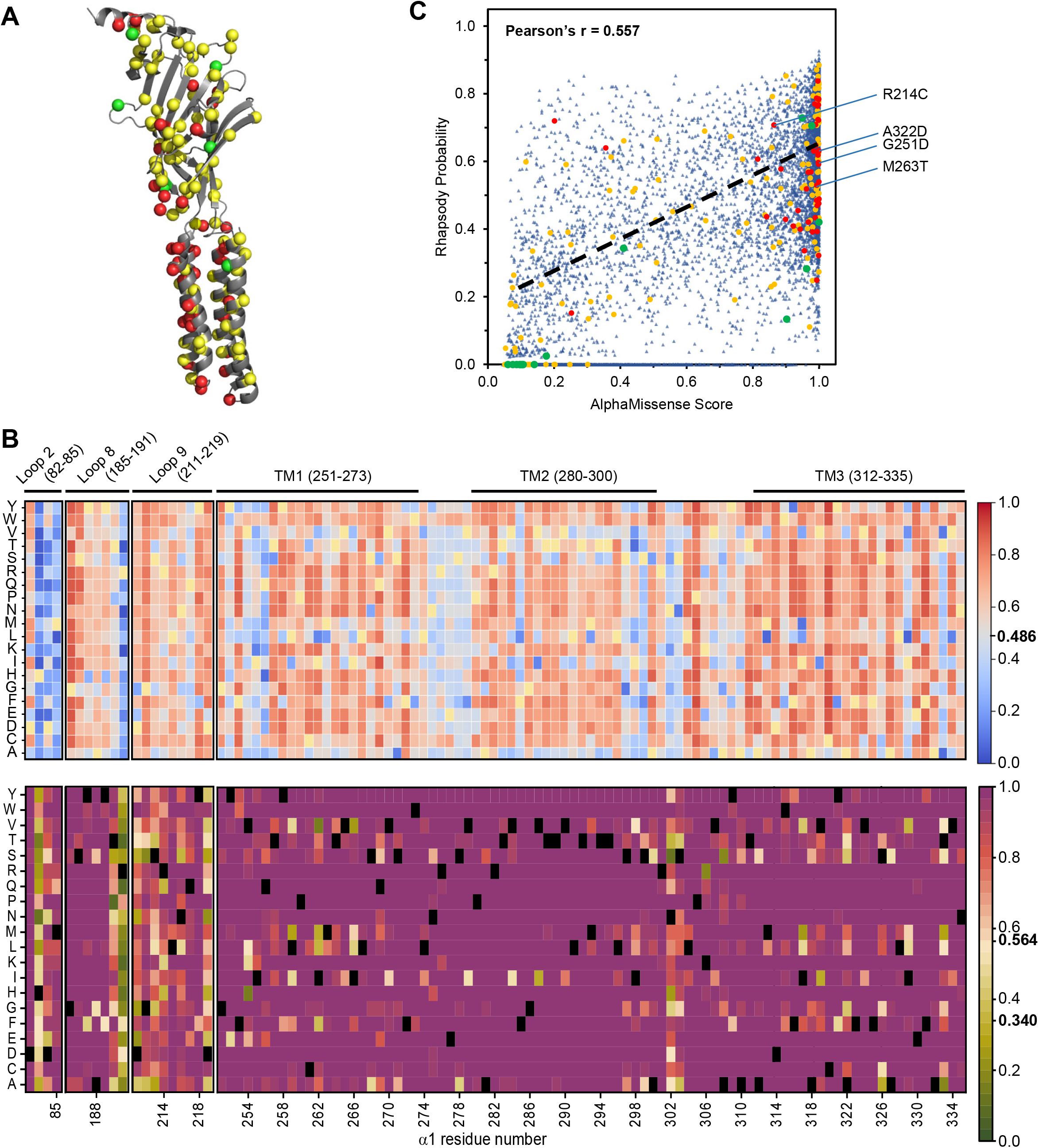
Pathogenicity prediction of GABA_A_ receptor α1 subunit, GABRA1. (**A**) Positions of clinical missense variants are displayed in the three-dimensional structure of α1 subunit, built from 6X3S.pdb using PyMOL. According to ClinVar annotation, Class I pathogenic variants are colored in red, Class II uncertain variants in yellow, and Class III benign variants in green. (**B**) Pathogenicity prediction of saturating substitutions of GABRA1. Heat maps display the pathogenic probability of selected key regions. The top picture is from Rhapsody (RS) prediction, whereas the bottom picture is from AlphaMissense (AM) prediction. Wild type amino acids were colored in yellow for RS, and black for AM. (**C**) Correlation between Rhapsody and AlphaMissense prediction. Red dots represent pathogenic variants, yellow dots represent uncertain variants, and green dots represent benign variants according to ClinVar annotation. If no predictions are available in Rhapsody, such variants are artificially assigned a value of zero for Rhapsody probability.

**Table 1.** The α1 subunit of GABA_A_R pathogenicity observations and predications.

| ClinVar Significance as Benign (Total 16)<sup>#</sup><br>AM as Benign 9, RS as Benign 5<sup>^</sup> |  |  |
| --- | --- | --- |
| Both AM RS as Benign |  | Both AM RS as Pathogenic |
| 1 |  | 2 |
| M108V |  | T66N I317T |

<sup>#</sup> Benign and Likely benign<sup>^</sup> Neutral and Probably neutral

| ClinVar Significance as Pathogenic (Total 48)<sup>\$</sup><br>AM as Pathogenic 42, RS as Pathogenic 29<sup>&</sup> | | | | | | |
| --- | --- | --- | --- | --- | --- | --- |
| Both AM RS as Pathogenic |  |  |  |  | Both AM RS as Benign |  |
| 28 |  |  |  |  | 1 |  |
| I45T | Y187D | M263K | T294I | **A322D** | R204C |  |
| D90H | A188D | **M263T** | L296F | F325L |  |  |
| D90N | **R214C** | L267F | S297R | A332V |  |  |
| D90V | R214S | P280L | K306T | N335H |  |  |
| F92S | N216I | T288I | A312T |  |  |  |
| P124L | **G251D** | V290M | D314N |  |  |  |

<sup>\$</sup> Pathogenic and Likely pathogenic<sup>&</sup> Deleterious and probably deleterious

| ClinVar Significance as Uncertain (Total 168)<sup>*</sup> |  |  |  |  |  |  |
| --- | --- | --- | --- | --- | --- | --- |
| Both AM RS as Pathogenic |  |  |  |  | Both AM RS as Benign |  |
| 58 |  |  |  |  | 18 |  |
| L46F | L111P | P167S | L267I | L328V | R64H | D226G |
| L49H | D125G | H178Q | V270G | F331I | V74I | Q231E |
| R56H | F127V | G185V | S271P | N335I | I89V | V287I |
| P59L | K144E | Y187C | S278Y | W344G | T109I | V349M |
| F73I | R147G | Y196H | A281E | S423L | H129Y | K410R |
| G78R | R147Q | G212R | T294R | F427S | R163K | T412S |
| D90G | R147W | Q217K | T295I | P428L | E201G | S425T |
| D90Y | I148F | Y218S | P305L | F431S | R204H | K418R |
| R94C | G152S | L220F | A308D | I433N | V207F | I433V |
| R94S | R159T | L221P | T311I | Y438C |  |  |
| W97R | T161K | T257I | V319M |  |  |  |
| D99G | C166W | C261R | Y321C |  |  |  |

<sup>\*</sup> Uncertain significance and Conflicting interpretations of pathogenicity

**Table 2.** The β2 subunit of GABA_A_R pathogenicity observations and predications.

| ClinVar Significance as Benign (Total 34)<sup>#</sup><br>AM as Benign 24, RS as Benign 5<sup>^</sup> |  |
| --- | --- |
| Both AM RS as Benign | Both AM RS as Pathogenic |
| 1 | 0 |
| F501I |  |

<sup>#</sup> Benign and Likely benign<sup>^</sup> Neutral and Probably neutral

| ClinVar Significance as Pathogenic (Total 39)<sup>\$</sup><br>AM as Pathogenic 39, RS as Pathogenic 25<sup>&</sup> | |
| --- | --- |
| Both AM RS as Pathogenic | Both AM RS as Benign |
| 25 | 0 |
| D125N P252L T286I I299F Y301C<br>Y244H P252T T287P I299S K303E<br>F245L P252S I288T P300L K303R<br>T249R L277S R293Q Y301H K303N<br>P252A V282A R293W Y301F A304V |  |

<sup>\$</sup> Pathogenic and Likely pathogenic<sup>&</sup> Deleterious and probably deleterious

| ClinVar Significance as Uncertain (Total 141)<sup>*</sup> |  |
| --- | --- |
| Both AM RS as Pathogenic | Both AM RS as Benign |
| 34 | 20 |
| L51M R141H R240S T280A A304T<br>G57S R141C G243S T281S I305T<br>I66T Y150H F245C T281A A319V<br>D80H M162R T249A V282F F331S<br>N109K Y183H I266M L283P R489W<br>W116C P230H A272V T284A R492H<br>N124S L234F A272T T286N | N32S R193H<br>P58S D195N<br>P58L N197K<br>P58A T203M<br>A61S K220E<br>I66V K221N<br>I101V K221R<br>P102S I504V<br>V148I V505I<br>E189V V511M |

<sup>\*</sup> Uncertain significance and Conflicting interpretations of pathogenicity

**Table 3.**
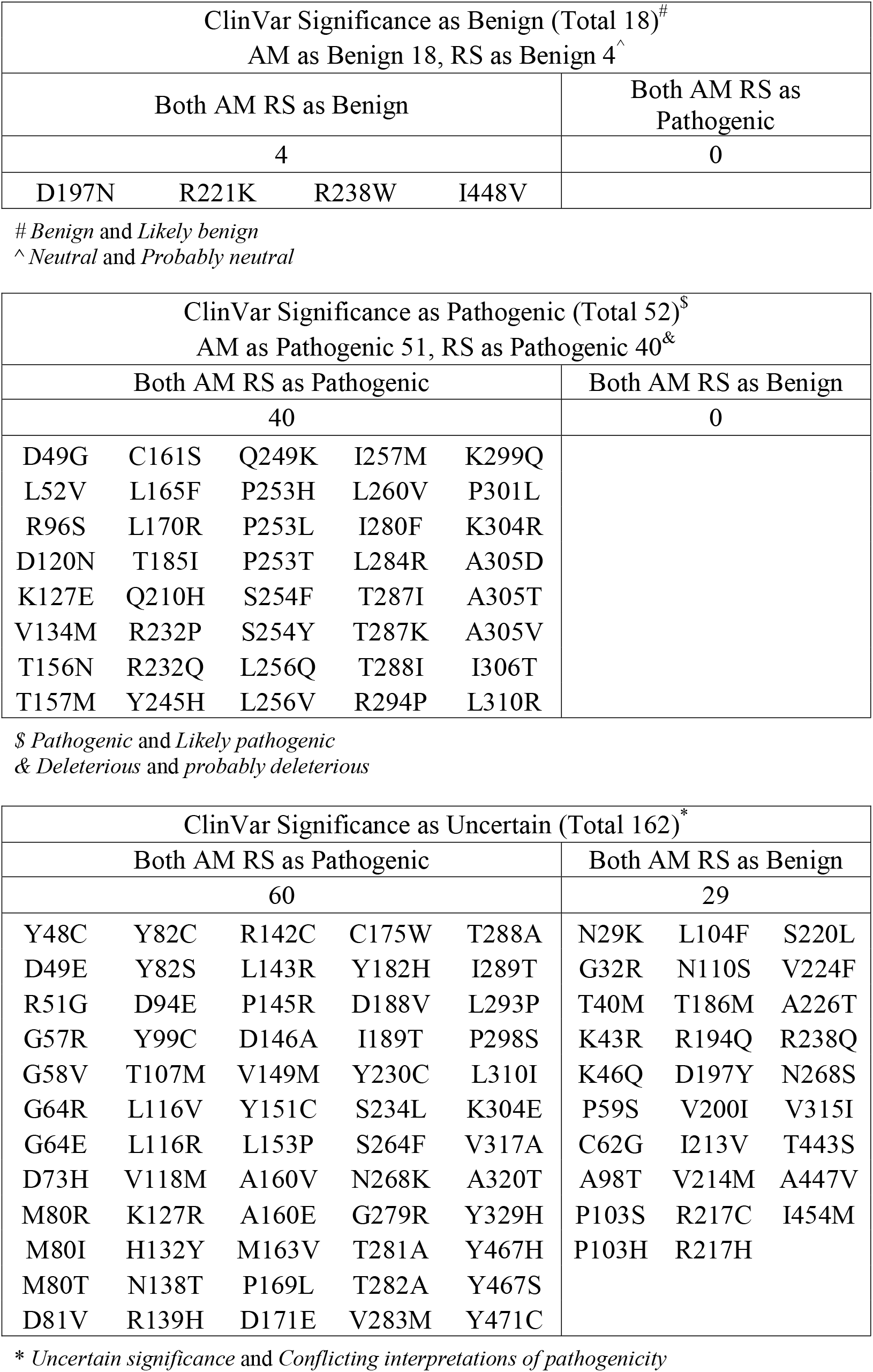
The β3 subunit of GABA_A_R pathogenicity observations and predications.

**Table 4.**
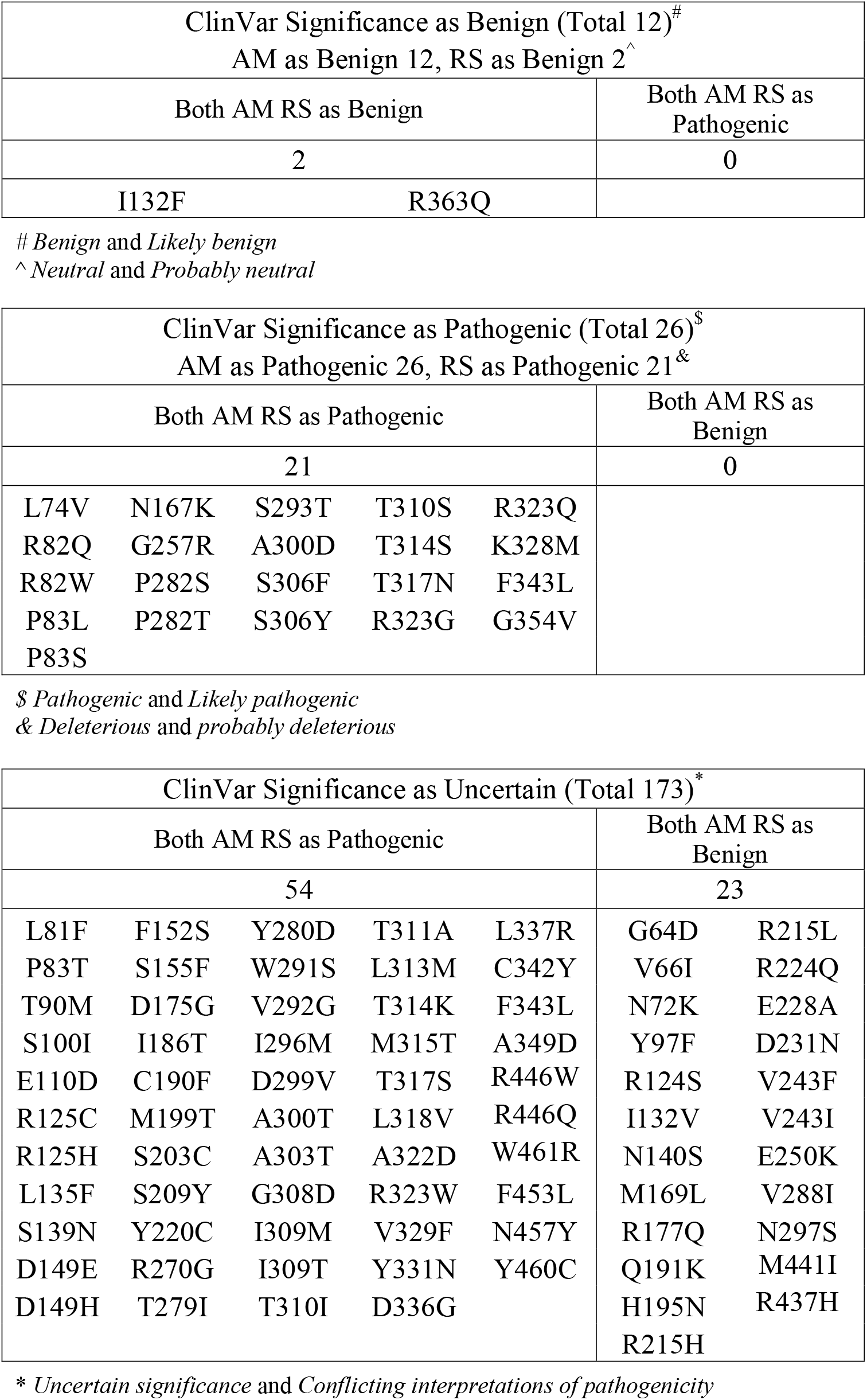
The γ2 subunit of GABA_A_R pathogenicity observations and predications.

The AM capacity enables the pathogenicity prediction for saturating substitutions in GABA_A_ receptor subunits. The saturating prediction provides information for missense variants that could be identified in clinic in the future as well as identifying critical regions in the protein sequences that are intolerant to variations. For example, transmembrane helices (TM1, TM2, and TM3) and loops connecting them, loop 2 connecting β2 sheet and β1 sheet, loop 8 (aka Loop B) connecting β8 sheet and β7 sheet, and loop 9 connecting β9 sheet and β8 sheet harbor many pathogenic clinical variants (**Figure 1D**). Consistently, AM prediction demonstrated that most of the variations in these positions resulted in high pathogenic probability (see bottom panels in **Figure 2B** for GABRA1, **Figure 3B** for GABRB2, **Figure 4B** for GABRB3, and **Figure 5B** for GABRG2), suggesting that it is critical to maintain the structural and functional integrity in these regions.

**Figure 3.**
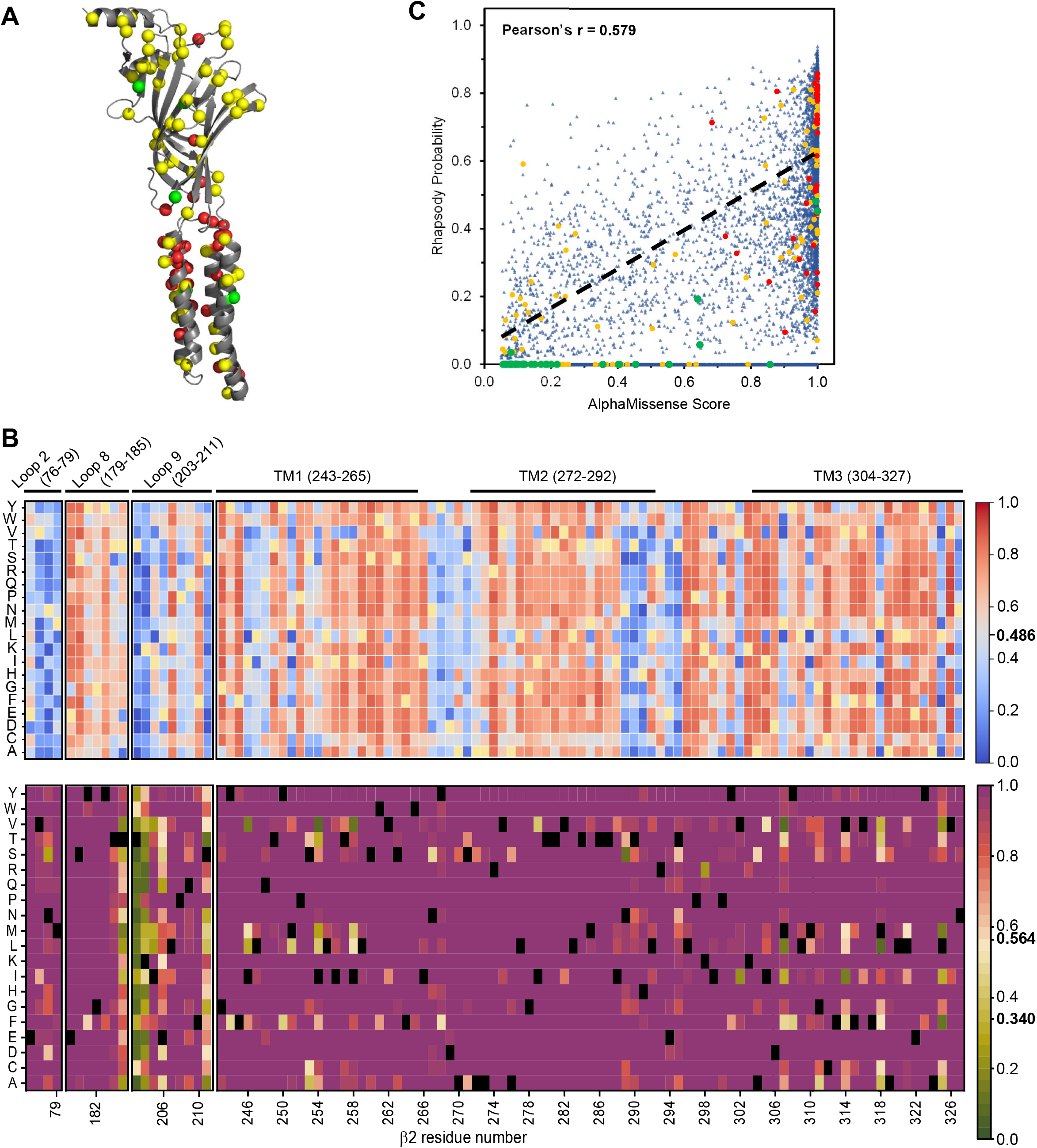
Pathogenicity prediction of GABA_A_ receptor β2 subunit, GABRB2. (**A**) Positions of clinical missense variants are displayed in the three-dimensional structure of β2 subunit, built from 6X3S.pdb using PyMOL. According to ClinVar annotation, Class I pathogenic variants are colored in red, Class II uncertain variants in yellow, and Class III benign variants in green. (**B**) Pathogenicity prediction of saturating substitutions of GABRB2. Heat maps display the pathogenic probability of selected key regions. The top picture is from Rhapsody (RS) prediction, whereas the bottom picture is from AlphaMissense (AM) prediction. Wild type amino acids were colored in yellow for RS, and black for AM. (**C**) Correlation between Rhapsody and AlphaMissense prediction. Red dots represent pathogenic variants, yellow dots represent uncertain variants, and green dots represent benign variants according to ClinVar annotation. If no predictions are available in Rhapsody, such variants are artificially assigned a value of zero for Rhapsody probability.

**Figure 4.**
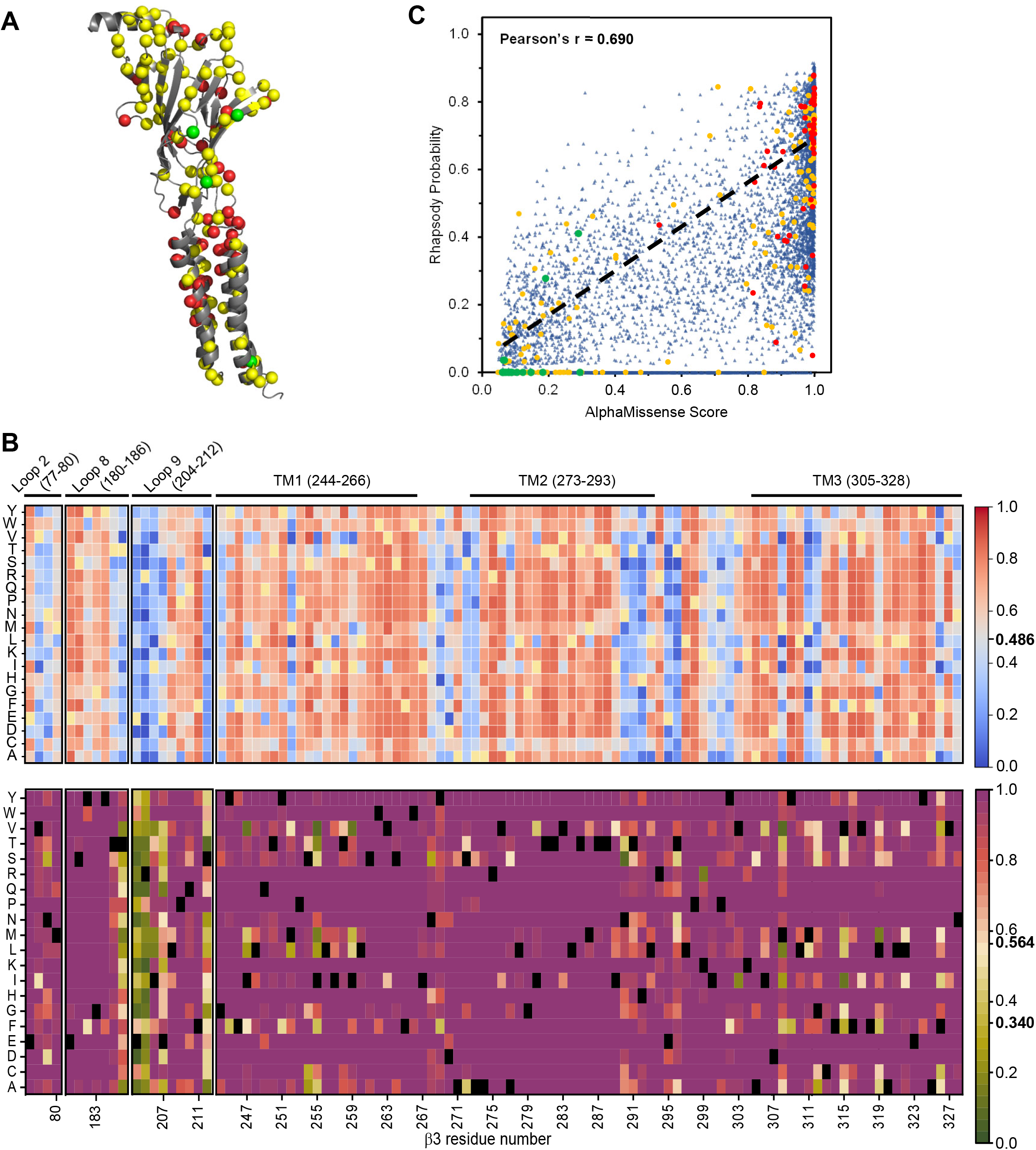
Pathogenicity prediction of GABA_A_ receptor β3 subunit, GABRB3. (**A**) Positions of clinical missense variants are displayed in the three-dimensional structure of β3 subunit, built from 6HUK.pdb using PyMOL. According to ClinVar annotation, Class I pathogenic variants are colored in red, Class II uncertain variants in yellow, and Class III benign variants in green. (**B**) Pathogenicity prediction of saturating substitutions of GABRB3. Heat maps display the pathogenic probability of selected key regions. The top picture is from Rhapsody (RS) prediction, whereas the bottom picture is from AlphaMissense (AM) prediction. Wild type amino acids were colored in yellow for RS, and black for AM. (**C**) Correlation between Rhapsody and AlphaMissense prediction. Red dots represent pathogenic variants, yellow dots represent uncertain variants, and green dots represent benign variants according to ClinVar annotation. If no predictions are available in Rhapsody, such variants are artificially assigned a value of zero for Rhapsody probability.

**Figure 5.**
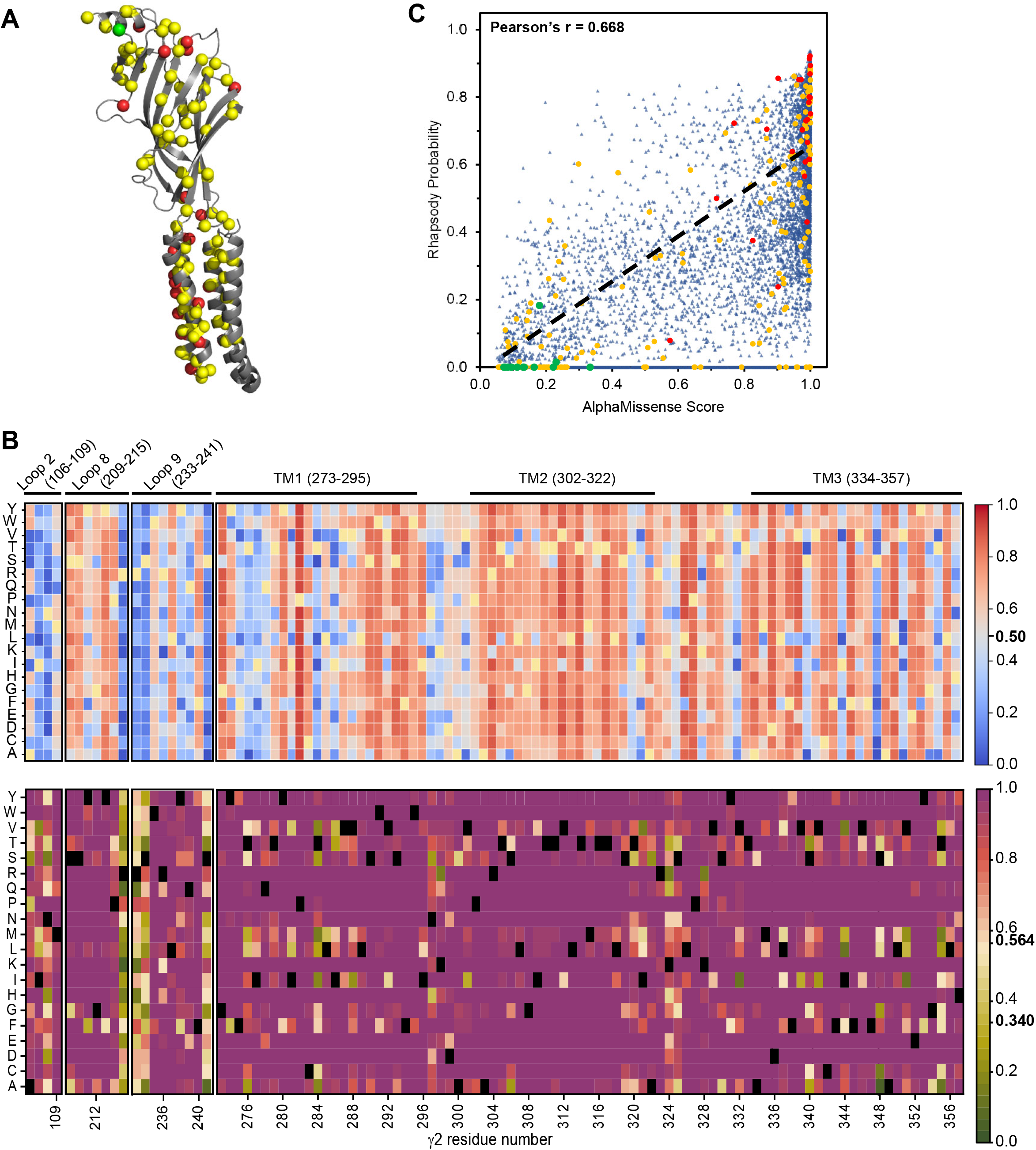
Pathogenicity prediction of GABA_A_ receptor γ2 subunit, GABRG2. (**A**) Positions of clinical missense variants are displayed in the three-dimensional structure of γ2 subunit, built from 6X3S.pdb using PyMOL. According to ClinVar annotation, Class I pathogenic variants are colored in red, Class II uncertain variants in yellow, and Class III benign variants in green. (**B**) Pathogenicity prediction of saturating substitutions of GABRG2. Heat maps display the pathogenic probability of selected key regions. The top picture is from Rhapsody (RS) prediction, whereas the bottom picture is from AlphaMissense (AM) prediction. Wild type amino acids were colored in yellow for RS, and black for AM. (**C**) Correlation between Rhapsody and AlphaMissense prediction. Red dots represent pathogenic variants, yellow dots represent uncertain variants, and green dots represent benign variants according to ClinVar annotation. If no predictions are available in Rhapsody, such variants are artificially assigned a value of zero for Rhapsody probability.

### Rhapsody (RS) pathogenicity correlates modestly with AlphaMissense predications

Rhapsody (RS) makes pathogenicity predictions by including structure-and dynamics-based information and is well performed for human missense variants ^[16]^. Previously, we applied RS to predict pathogenicity of GABRA1 variants, and several variants were further verified experimentally ^[13b]^. Here, we used RS for the saturating pathogenicity predictions and compared the results with AM predictions. Since RS relies on the known cryo-EM structures of GABA_A_ receptors, the RS predictions are not available for the positions that do not have such structural information, such as those in the signal peptide and most of the long TM3-4 loop. We evaluated the correlation between RS and AM using a Pearson’s coefficient from linear regression fitting of values of RS probability vs AM score. RS pathogenicity has a modest correlation with AM prediction with a Pearson’s coefficient of 0.557 for GABRA1 (**Figure 2C**), 0.579 for GABRB2 (**Figure 3C**), 0.690 for GABRB3 (**Figure 4C**), and 0.668 for GABRG2 (**Figure 5C**). In addition, similar to AM prediction, RS also revealed that several regions in the primary protein sequences, such as TM1-TM3 and their connecting loops, and loop 2, loop 8, and loop 9 play an essential role in the maintenance of GABA_A_ receptor function (see top panels in **Figure 2B** for GABRA1, **Figure 3B** for GABRB2, **Figure 4B** for GABRB3, and **Figure 5B** for GABRG2).

#### AlphaMissense and Rhapsody provide guidance for the investigation of missense variants

Pathogenic annotations and predications have highest consistency among CV, AM, and RS. For GABRA1, CV annotated 48 Class I pathogenic variants, among which AM predicts 42 and RS predicts 29; both AM and RS predicted 28 common variants to be pathogenic (**Table 1**). Previously, we showed that R214C, G251D, M263T and A322D in GABRA1 impaired proteostasis and reduced current amplitudes ^[13b]^. For GABRB2, CV annotated 39 Class I pathogenic variants, among which AM predicts 39 and RS predicts 25; both AM and RS predicted 25 common variants to be pathogenic (**Table 2**). Previously, it was reported that P252L and V282A in GABRB2 resulted in reduced current amplitudes ^[23]^. For GABRB3, CV annotated 52 Class I pathogenic variants, among which AM predicts 51 and RS predicts 40; both AM and RS predicted 40 common variants to be pathogenic (**Table 3**). Previously, it was reported that a number of variants in GABRB3 caused loss-of-function or gain-of-function defects ^[24]^. For GABRG2, CV annotated 26 Class I pathogenic variants, among which AM predicts 26 and RS predicts 21; both AM and RS predicted 21 common variants to be pathogenic (**Table 4**). Previously, it was reported that several variants in GABRG2, including P282S, R323Q, and F343L, reduced cell surface expressions and current amplitudes ^[25]^.

Since ClinVar is inadequate in annotating clinical variants (∼72% of missense GABA_A_ variants in ClinVar have uncertain clinical significance), AM and RS provide guidance to fill the gap for the interpretation of missense variants. For GABRA1, CV had 168 Class II uncertain variants with clinical significance as uncertain or conflicting interpretations of pathogenicity, among which both AM and RS predicted 58 common variants to be pathogenic and 18 common variants to be benign (**Table 1**). For GABRB2, CV had 141 Class II uncertain variants, among which both AM and RS predicted 34 common variants to be pathogenic and 20 common variants to be benign (**Table 2**). For GABRB3, CV had 162 Class II uncertain variants, among which both AM and RS predicted 60 common variants to be pathogenic and 29 common variants to be benign (**Table 3**). For GABRG2, CV had 173 Class II uncertain variants, among which both AM and RS predicted 54 common variants to be pathogenic and 23 common variants to be benign (**Table 4**). The differentiation of the CV uncertain variants by AM and RS provides valuable information for further validation and interpretation.

## CONCLUSION

Advances in genome sequencing have identified a growing number of variants associated with a broad spectrum of epilepsy ^[26]^. Currently, over 1000 clinical variants have been identified in genes encoding GABA_A_ receptor subunits ^[27]^. However, a substantial knowledge gap exists for the interpretation of the clinical significance of these variants. Recent development of computational approaches such as machine learning enables the prediction of the pathogenicity of genetic variants ^[28]^. We utilized two state-of-the-art modeling approaches, namely AlphaMissense and Rhapsody, to perform the saturating substitution analysis for the major subunits of GABA_A_ receptors. AlphaMissense and Rhapsody predictions correlated modestly with each other. The predicted pathogenicity agreed best with the ClinVar annotated pathogenic variants. Furthermore, the prediction differentiated the ClinVar uncertain variants to different classes, giving directions for future experiments and interpretations using high-throughput approaches. Since each GABA_A_ variant is rare, the saturating substitution prediction provides a valuable clue about the functional consequence of the rare variant and contributes to a tailored treatment option. It is likely that proteostasis deficiency remains the major disease-causing mechanism for pathogenic GABA_A_ receptor variants after a comprehensive characterization. As such, adapting the proteostasis network using proteostasis regulators has the promise to rescue the function of a variety of variants since this strategy targets the global cellular signaling pathways ^[14b]^. In addition, pharmacological chaperones that directly bind GABA_A_ receptors can stabilize the variants to restore their trafficking and function in a receptor-specific way ^[13b]^. Both proteostasis regulators and pharmacological chaperones have the promise to be further developed to treat genetic epilepsy resulting from GABA_A_ receptor dysfunction. Since proteostasis regulators and pharmacological chaperones have distinct mechanism of action, their combination use is expected to lead to additive or synergistic effects on correcting the function of pathogenic GABA_A_ receptor variants.

## METHODS

### Source data collection from AlphaMissense (AM), ClinVar (CV) and Rhapsody (RS)

AlphaMissense (AM) predictions were downloaded for all single amino acid substitutions in the human proteome data ^[15]^, and further extracted into separate files after searching for the GABA_A_ receptor subunits with the following accession numbers (Uniprot IDs): P14867 for α1 (gene name *GABRA1*), P47870 for β2 (gene name *GABRB2*), P28472 for β3 (gene name *GABRB3*), and P18507 for γ2 (gene name *GABRG2*). The total numbers of 8664, 9728, 8987, and 8873 values were available for GABRA1, GABRB2, GAGRB3, and GABRG2, respectively.

ClinVar (CV) data was downloaded from the CV website (https://www.ncbi.nlm.nih.gov/clinvar) after searching for GABRA1, GABRB2, GAGRB3, and GABRG2, and was further filtered for missense variants by removing frameshift, nonsense, splice site et al. Due to variable splicing isoforms used in CV datasets, certain variants were manually checked to match AM amino acid residue numbers. Filtering yielded 232, 214, 232, and 211 missense variant pathogenicity annotations for GABRA1, GABRB2, GAGRB3, and GABRG2, respectively. Moreover, the missense variants were categorized into the following classes for further evaluation: i) pathogenic, including pathogenic and likely pathogenic, equivalent to the AM predications of pathogenic; ii) uncertain, including uncertain significance and conflicting interpretations, equivalent to the AM predications of ambiguous; iii) benign, including benign and likely benign, equivalent to the AM predications of benign.

Rhapsody (http://rhapsody.csb.pitt.edu/) was used to run *in silico* saturation mutagenesis analysis for GABRA1, GABRB2, GAGRB3 and GABRG2 using their Uniprot IDs ^[16]^. The results were downloaded. Probability scores and classifications were utilized for further analysis and comparison. Particularly, the equivalent categories were as the following: i) pathogenic, including deleterious and likely deleterious, equivalent to the AM predications of pathogenic; ii) benign, including neutral and likely neutral, equivalent to the AM predications of benign. The cut-off values to differentiate the two categories of pathogenic and benign are 0.486 for GABRA1, GABRB2 and GABRB3, and 0.50 for GABRG2. There was unavailable output for a number of variants that were excluded, such as those in the signal peptide regions and in the regions between TM3 and TM4, which had unavailable structural information. Therefore, the total numbers of 6669, 6194, 6593, and 6289 values were available for GABRA1, GABRB2, GAGRB3, and GABRG2, respectively.

### Analysis

Sequence alignment for human GABRA1 (Uniprot #: P14867), GABRB2 (Uniprot #: P47870), GABRB3 (Uniprot #: P28472), and GABRG2 (Uniprot #: P18507) was carried out using Clustal Omega, a multiple sequence alignment program ^[29]^. Structural illustration of GABA_A_ receptors was generated with the PyMOL 2.5 software using 6X3S.pdb for α1β2γ2 receptors and 6HUK.pdb for α1β3γ2 receptors ^[11a, 12]^. Heatmaps were generated using the Origin 2023 software. Wild type amino acids were colored in yellow for Rhapsody, and black for AlphaMissense. Pearson’s coefficients from linear regression and line fitting were generated using the Origin 2023 software as well.

## Supplementary information

Supplementary information includes one supplemental figure and four supplemental tables.

## Acknowledgements

This work was supported by the National Institutes of Health (R01NS105789 and R01NS117176 to TM). We congratulate Dr. Jeffery Kelly for the receipt of the 2023 Wolf Prize in Chemistry.

## Author Contributions

Conceptualization, YW and TM; Data collection and analysis, YW, GV, and TM; Funding acquisition, TM; Writing, original draft, YW and TM; Writing, review and editing, YW, GV, and TM. All authors have read and agreed to the final version of the manuscript.

---

**Figure S1:**
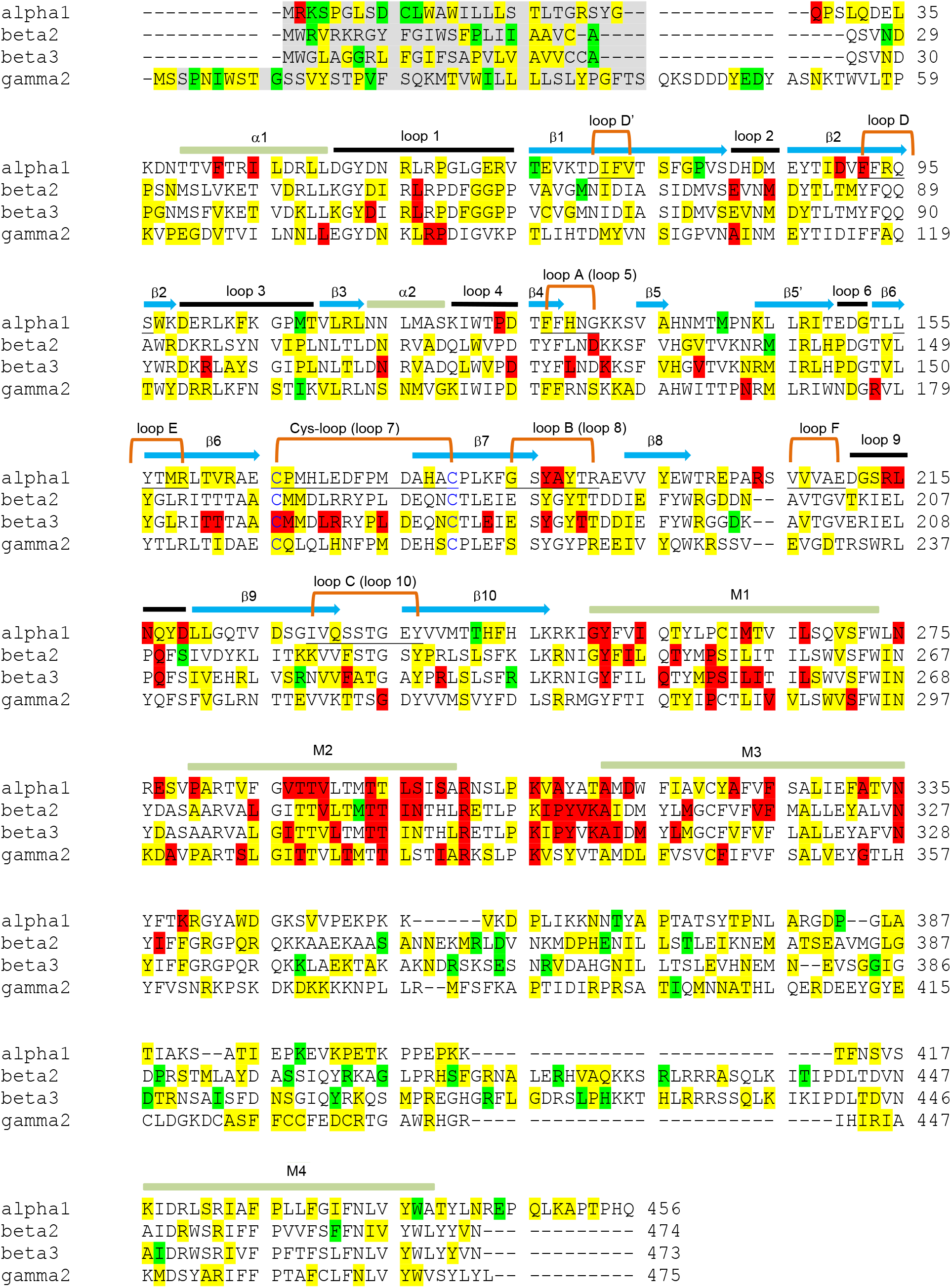
Primary sequence alignment of the major subunits of GABA_A_R: alpha1 (gene name *GABRA1*), beta2 (gene name *GABRB2*), beta3 (gene name *GABRB3*), gamma2 (gene name *GABRG2*). Amino acids of interest are colored according to *ClinVar* clinical significance (downloaded on 10/25/2023), red: pathogenic/likely pathogenic; yellow: uncertain/conflicting interpretations; green: benign/likely benign.

## Notes

### Competing Interest Statement

The authors have declared no competing interest.

